# ELONGATED HYPOCOTYL 5 regulates light-mediated squalene biosynthesis and development in *Arabidopsis thaliana*

**DOI:** 10.1101/2024.12.20.629791

**Authors:** Pranshu Kumar Pathak, Aruba Khan, Ashish Sharma, Nivedita Singh, Gurpreet Sandhu, Prabodh Kumar Trivedi

## Abstract

Terpenoids are diverse groups of metabolite families that are crucial for plant development and required in cosmetics and pharmacological industries. Various development processes and environmental factors, including light, have been shown to affect terpenoid biosynthesis. However, regulatory factors involved in such regulation have not been explored much. Squalene synthases (SQSs), key enzymes in the terpenoid pathway, are pivotal for sterol and triterpene biosynthesis across various organisms. In this study, we report that *AtSQS1* expression and squalene content are higher in the dark, and light through ELONGATED HYPOCOTYL 5 (HY5) negatively regulates the *AtSQS1* gene and squalene biosynthesis in *Arabidopsis thaliana*. Our study shows that the *AtSQS1* gene is unaffected in the *hy5-215* mutant during light and dark conditions, but it is down-regulated in WT and HY5OX lines. The histochemical GUS assays and GFP expression patterns indicate a negative regulation of squalene biosynthesis by AtHY5. Yeast one-hybrid assays, EMSA, and ChIP experiments have confirmed the physical binding of AtHY5 on light-responsive elements of the *AtSQS1* promoter. We have validated our results by developing *AtSQS1* promoter: reporter transgenic lines in WT, *hy5-215* mutants, and HY5OX backgrounds and conducted expression, metabolite quantification, and histochemical GUS assays. The metabolites quantification of squalene and phytosterols through GC-MS further confirms that AtHY5 negatively regulates the squalene biosynthesis in a light-dependent manner in Arabidopsis.

## Introduction

Plants synthesize an extensive array of compounds crucial for their growth and development, encompassing vital processes such as seed germination, reproduction, and photosynthesis, collectively termed primary metabolites. Beyond these fundamental functions, plants also synthesise a rich spectrum of bioactive compounds referred to as secondary metabolites. Unlike primary metabolites, these secondary compounds are not directly tied to essential plant processes; they play pivotal roles in mitigating diverse biotic and abiotic stressors (Al-Khayri et al. 2023; Gautam et al. 2023; Sharma et al. 2023; Sinha et al. 2024). Among these compounds, terpenoids are natural products based on isoprene and play essential roles in the metabolism of all living organisms (Nagegowda et al. 2020; Câmara et al. 2024). The formation of terpenoids typically involves continuous head-to-tail addition of building blocks, namely isoprene diphosphate (IPP) and its isomer dimethylallyl diphosphate (DMAPP) (Vranová et al. 2013; Sonawane et al. 2016). Initially, a head-to-tail condensation of IPP and DMAPP results in the production of geranyl diphosphate (GPP), serving as the precursor of monoterpenoids (**Fig. 1A**). Subsequent addition of IPP leads to the formation of farnesyl diphosphate (FPP) and geranylgeranyl diphosphate (GGPP), the precursors of sesquiterpenoids and diterpenoids, respectively. Additionally, two molecules of FPP and GGPP are condensed head-to-head to create squalene and phytoene, which act as the precursors of triterpenoids and tetraterpenoids. These linearized precursors undergo cyclization and post-modifications (e.g., oxidation, acetylation) to generate various terpenoids (Gupta et al. 2015; Câmara et al. 2024).

**Figure 1:**
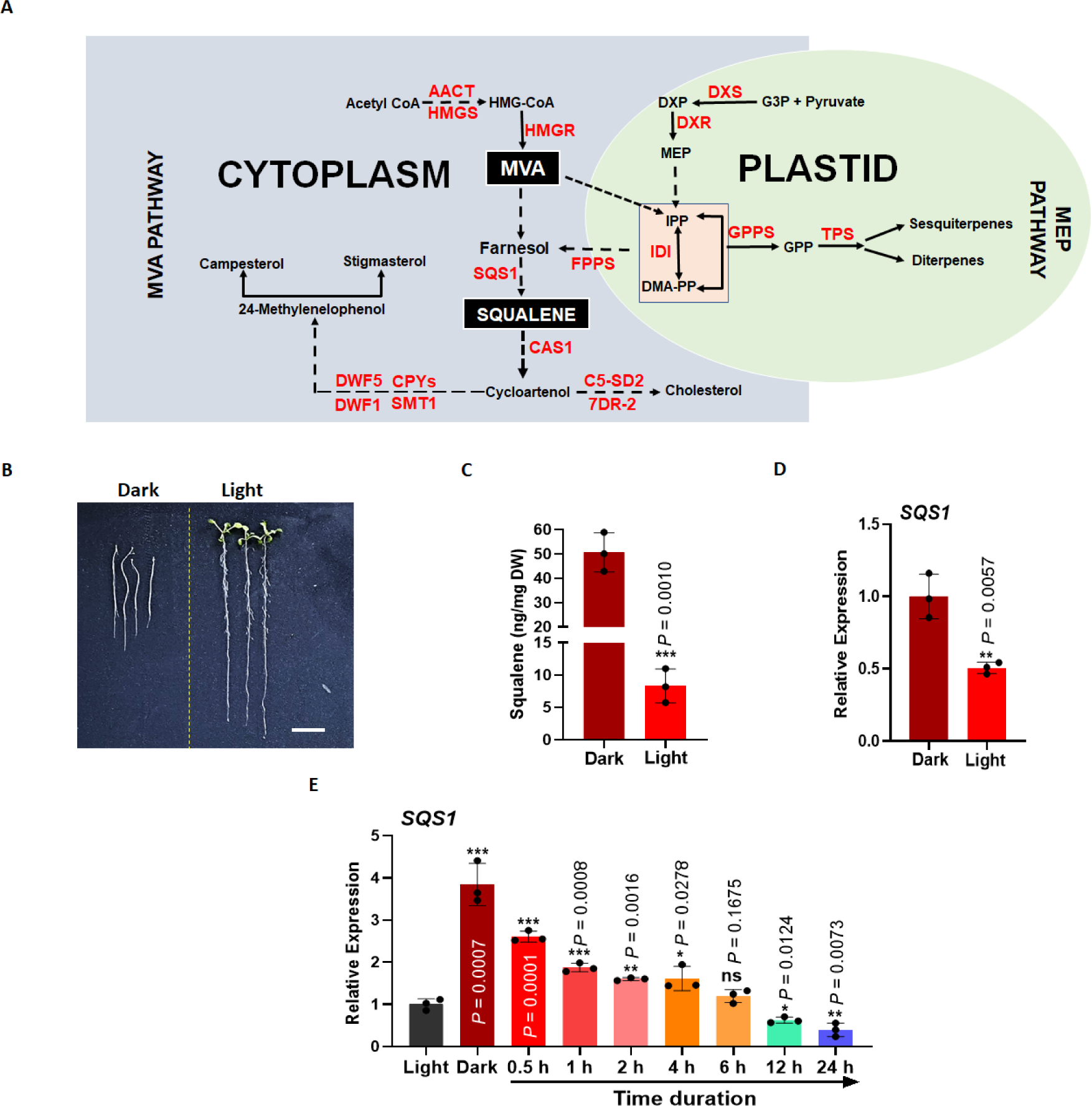
Light negatively regulates squalene biosynthesis in *Arabidopsis thaliana*. **(A)** Schematic representation of the biosynthesis pathway of terpenoids involves both the mevalonate (MVA) and methylerythritol phosphate (MEP) pathways. **(B)** Ten-day-old Arabidopsis seedlings grown in light and dark conditions. (**C**) Quantification Squalene content in light- and dark-grown 10-day-old Arabidopsis WT seedlings through GC-MS. **(D)** RT-qPCR expression analysis of terpenoid biosynthetic structural gene, *AtSQS1,* in light- and dark-grown 10-day-old WT Arabidopsis seedlings. *Tubulin* was used as the endogenous control to normalize the relative expression levels. **(E)** Expression analysis of *AtSQS1* in WT seedlings grown for 10 days in the dark followed by exposure to white light for different time intervals. The experiment was conducted independently at least three times, yielding similar results each time. The statistical analysis was performed using two-tailed Student’s t-tests. The data are plotted as means ± s.d. The error bars represent standard deviations. The asterisks indicate significant differences: *P< 0.001.

In general, the MEP pathway is the source of C5 prenyldiphosphates necessary for synthesizing C10 monoterpenes, C20 diterpenes, and C40 tetraterpenes. Conversely, the MVA pathway provides the common precursors for the synthesis of C15 sesquiterpenes, C27-29 sterols, and C30 triterpenes. Squalene synthases (SQSs) are essential for the condensation of two farnesyl diphosphate (farnesyl-PP) molecules, producing squalene, a pivotal precursor in sterol biosynthesis (Santana et al. 2020). Operating as a membrane-bound enzyme, SQSs are implicated in phytosterol biosynthesis, contributing to membrane interactions and influencing the biological activities of diverse cell proteins engaged in signal transduction, membrane formation, and the regulation of cell growth (Liao et al. 2018; Fu et al. 2023; Zhang et al. 2023). In *Arabidopsis thaliana* possesses two SQSs-annotated genomic sequences, At4g34640 (SQS1) and At4g34650 (SQS2), arranged in a tandem array. Out of these two isoforms, only SQS1 is capable of coding for a functional protein, whereas SQS2 is regulated solely at the transcriptional level (Busquets, et al. 2008).

Light plays a significant role in the growth and development of plants. Plants undergo skotomorphogenesis in the absence of light, whereas seedlings exhibit photomorphogenesis in the presence of light (Sullivan and Deng, 2003; Tripathi et al. 2019). During the dark-to-light transition, light-activated photoreceptors coordinate, and photomorphogenesis by transcriptional regulation (Huang et al. 2014; Podolec and Ulm, 2018; Kreiss et al. 2023). ELONGATED HYPOCOTYL5 (HY5) operates downstream of all photoreceptors and serves as a central regulator in the light signal transduction pathway (Ponnu and Hoecker, 2022; Mankotia et al. 2024). The abundance of HY5 is meticulously regulated by CONSTITUTIVELY PHOTOMORPHOGENIC/DEETIOLATED/FUSCA (COP/DET/FUS) proteins, whose biochemical activities are suppressed by photoreceptors upon light exposure (Holtkotte et al. 2017; Podolec and Ulm, 2018). As a b-ZIP-type transcription factor, HY5 selectively binds to 5’-NNACGTNN-3’-containing (ACE) *cis*-elements found in the promoters of about one-third of Arabidopsis genes, thereby regulating their transcription and ultimately promoting photomorphogenic development under light conditions (Lee et al. 2007; Zhang et al. 2011). In the darkness, HY5 undergoes ubiquitination and degradation facilitated by COP1, whereas its abundance increases significantly in light conditions. These components, in collaboration with COP1 and HY5, work synergistically to precisely regulate light-mediated different processes in plants (Huang et al. 2014; Hoecker, 2017; Podolec and Ulm, 2018; Singh et al. 2024; Sinha et al. 2024; Gaddam et al. 2024).

There are a few reports of the influence of light as well as light signalling components on the terpenoid biosynthesis pathway (Kawoosa et al. 2010; Micheal et al. 2020; He et al. 2021; Liu et al. 2023; Micheal et al. 2024). Regulation of terpenoid biosynthesis by other transcription factors like MYB (Han et al. 2022) and WRKY (Fu et al. 2021) is also induced by light (Singh et al. 2024). Light plays a critical role in regulating terpenoid production in plants as it affects plant organs. There are reports about HY5 being a central regulator of light signal transduction in controlling the transcriptional regulation of terpenoid biosynthesis under broad spectra i.e., UV, Red, and Blue light (Aviles et al. 2024). However, the mechanism and role of light-associated components in phytosterol biosynthesis have not been thoroughly investigated. Our study indicates the presence of light-dependent transcriptional regulation in the biosynthesis of terpenoids (specifically, squalene biosynthesis) and phytosterols in *A. thaliana*. Physiological, biochemical, and molecular analyses using different tools suggest a light-dependent regulatory pattern controlled by HY5. We used *hy5-215* mutant, AtHY5 overexpressing (AtHY5OX), and AtSQS1-overexpression lines in different backgrounds to validate that AtHY5 negatively regulates squalene biosynthesis by regulating *AtSQS1* expression via interaction with *cis*-regulatory elements.

## Results

### Squalene biosynthesis is regulated by light

The terpenoid biosynthesis involves both the MVA and MEP pathways (**Fig. 1A**), and some of the genes involved in these pathways are known to be regulated by light through HY5 **(Supplemental Fig. S1)**. To investigate whether squalene biosynthesis and *AtSQS1* are regulated by light, wild-type (WT) seedlings were grown in the light and dark for 10 days and analysed for transcript levels of *AtSQS1* and squalene content. The phenotypic differences showed a clear difference between skotomorphogenesis and photomorphogenesis in 10-day-old light and dark-grown seedlings (**Fig. 1B**). A significantly elevated level of squalene was observed in dark-grown seedlings compared to light-grown seedlings, suggesting squalene accumulation might be negatively regulated by the light (**Fig. 1C**). The transcript level of *AtSQS1* was also significantly lower in light-grown seedlings compared to dark-drown seedlings (**Fig. 1D**). To study the light-responsiveness of *AtSQS1*, 10-day-old dark-grown seedlings were exposed to white light for different time intervals, and the expression of *AtSQS1* was analysed. The expression of *AtSQS1* decreased in a time-dependent manner after light exposure (**Fig. 1E**). To study the possible mechanism for the light-dependent regulation of *AtSQS1*, the promoter of the gene was analysed. The analysis suggested the presence of potential light-responsive elements (LRE) in the *AtSQS1* promoter (**Supplemental Table S1**). All these results together suggest a possible role of light-associated transcription factors in the regulation of expression and squalene biosynthesis.

### Involvement of HY5 in light-dependent expression of *AtSQS1*

Previous findings suggested transcription factor HY5 as a pivotal regulator of light-regulated processes in plants, impacting various aspects such as cell elongation, proliferation, and the accumulation of secondary metabolites (Gaddam et al. 2024; Singh et al. 2024). A previous study utilizing genome-wide ChIP experiments revealed that HY5 influences the expression of approximately one-third of Arabidopsis genes, with promoters of nearly 3000 genes directly binding to HY5 (Lee et al. 2007; Zhang et al. 2011). HY5 mutant (*hy5-215*) has long hypocotyl, whereas, on overexpression of HY5 (AtHY5OX), hypocotyl becomes short in light. In dark condition, long hypocotyl was observed in *hy5-215 and* AtHY5OX plants when compared to the WT **(Supplemental Fig. S2)**. To study whether light-dependent expression of *AtSQS1* and squalene synthesis is through HY5, the expression of *AtSQS1* and squalene and cholesterol content were analysed in 10-day-old seedlings of WT, *hy5-215,* and AtHY5OX plants. The analysis indicated enhanced expression of *AtSQS1* as well as increased squalene content in *hy5-215* mutant plants, whereas significantly lower expression and content were observed in AtHY5OX lines when compared to WT **(Fig. 2, A-C)**. Similarly, expression of *AtSQS1*, squalene, terpenoid, and cholesterol contents was studied in dark and light-grown seedlings of WT, *hy5-215*, and AtHY5OX plants. Analysis suggested that there was no significant difference in *AtSQS1* expression as well as squalene content in light-grown seedlings of mutants when compared to their dark-grown counterparts, whereas the terpenoid content was reduced in light conditions **(Fig. 2, D-F)**. On the other hand, the *AtSQS1* expression, squalene content, and terpenoid content were reduced substantially in light conditions compared to their dark counterparts **(Fig. 2, D-F)**.

**Figure 2:**
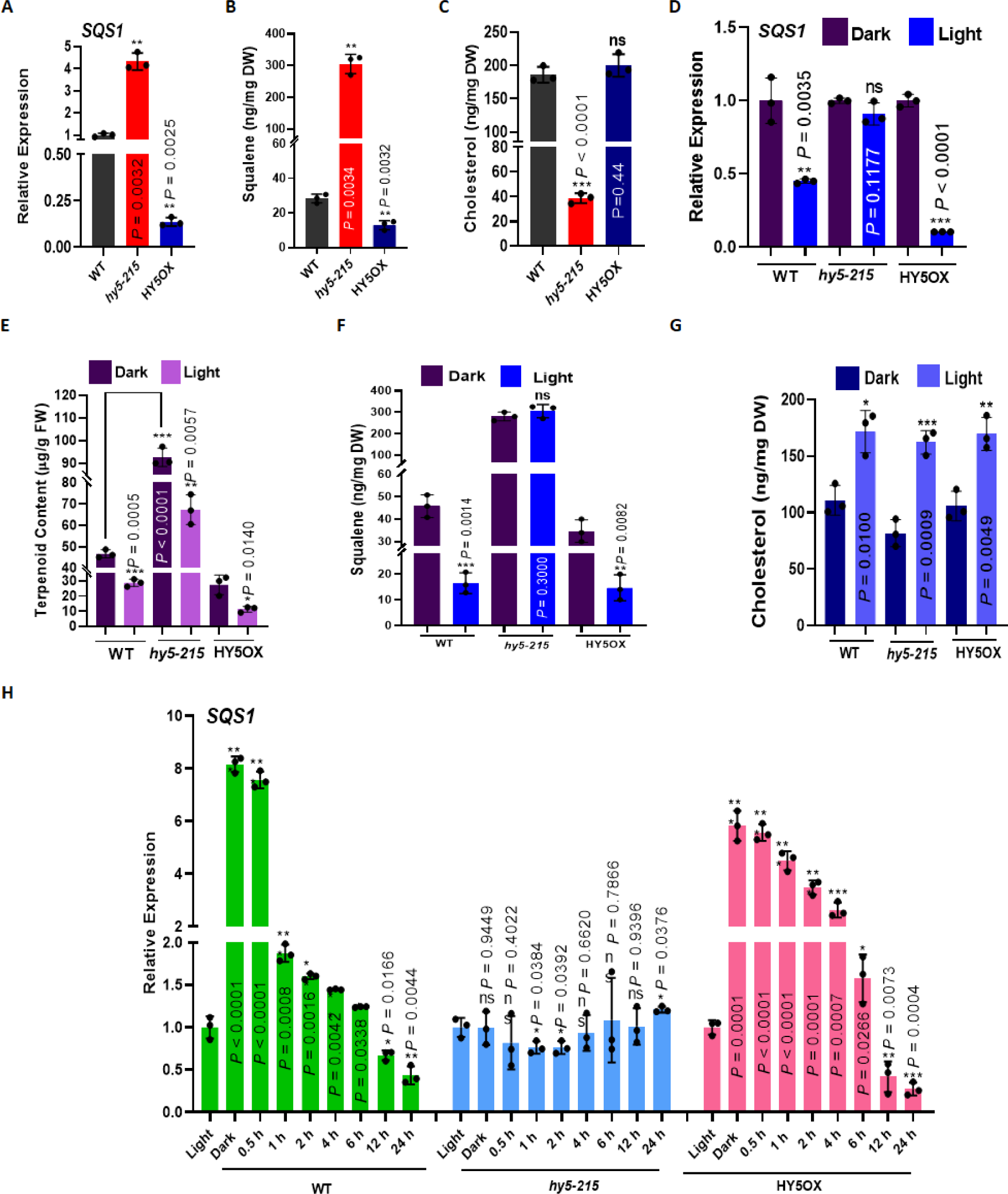
Light-dependent expression of *AtSQS1* is regulated by Elongated Hypocotyl 5 (HY5). **(A)** Expression analysis of terpenoid biosynthetic structural gene, *AtSQS1,* in 10-day-old seedlings of WT, *hy5-215*, and HY5OX Arabidopsis plants grown under light conditions. *Tubulin* was used as the endogenous control to normalize the relative expression levels (the small open circles represent the individual values). **(B and C)** Quantification of Squalene and Cholesterol content in the WT, *hy5-215*, and HY5OX seedlings were estimated in light conditions. **(D)** Expression analysis of terpenoid biosynthetic gene, *AtSQS1,* in 10-day-old WT, *hy5-215*, and HY5OX seedlings grown under light and dark conditions. *Tubulin* was used as the endogenous control to normalize the relative expression levels (the small open circles represent the individual values). **(E, F, and G)** Quantification of total terpenoid and squalene contents and cholesterol levels in the WT, *hy5-215*, and HY5OX seedlings grown under light and dark conditions. **(H)** Expression analysis of *AtSQS1* in WT, *hy5-215*, and HY5OX Arabidopsis seedlings grown for 10 days in the dark followed by exposure to white light for different time intervals (from dark to 24 h of white light). The experiment was conducted independently at least three times, yielding similar results each time. The statistical analysis was performed using two-tailed Student’s t-tests. The data are plotted as means ± s.d. The error bars represent standard deviations. The asterisks indicate significant differences: *P< 0.001.

As squalene content was enhanced in *hy5-215* plants, which led us to study the cholesterol content also in the same lines. The cholesterol content was significantly high in *hy5-215* plants, which might be due to increased squalene in these plants, as cholesterol is a downstream metabolite of the same pathway **(Fig. 2G)**. Dark-grown seedlings of WT, *hy5-215*, and AtHY5OX plants were exposed to light in a time-dependent manner; expression of *AtSQS1* decreased gradually with time in WT and AtHY5OX plants. However, *hy5-215* plants showed no influence of light exposure in the expression **(Fig. 2H)**. The relative expression of *AtCHS* and *AtHY5* genes was also analysed, and they gradually increased with light exposure as these genes are known to be positively regulated by light (Catalá et al. 2011; Gangappa and botto 2016) (**Supplemental Fig. S3, A and B**). The absence of light signalling components in mutant lines led to increased expression of *AtSQS1*, which clearly indicates AtHY5 as a negative regulator of *AtSQS1*. These findings suggested that AtSQS1 expression is modulated by light via transcription factor i.e., HY5.

### AtHY5 interacts with *AtSQS1* promoter

In previous experiments, we observed that *AtSQS1* expression decreases when dark-grown seedlings are exposed to light, suggesting that AtHY5 may act as a negative regulator of *AtSQS1*. To test this hypothesis, we conducted studies to investigate the interaction between the *AtSQS1* promoter and AtHY5. The ∼1.9 kb proximal promoter region of *AtSQS1* was analyzed for light-responsive *cis*-elements, suggesting the presence of such elements upstream of the transcription start site **(Fig. 3A, Supplemental Fig. S4A, Supplemental Table S1)**. A yeast one-hybrid (Y1H) assay was performed to confirm the physical interaction between AtHY5 and the *AtSQS1* promoter. The promoter region was cloned into the pAbAi vector to create the bait construct (ProAtSQS1-pAbAi), which was validated for AbA**^r^** resistance on-Ura medium with AbA**^r^** concentrations ranging from 100 to 300 ng/ml **(Supplemental Fig. S4B)**. The prey construct was prepared by cloning AtHY5 into the pGADT7-AD vector. Co-transformation of both constructs into yeast cells followed by growth on SD/-Leu/AbA selection medium confirmed the interaction through the presence of positive colonies **(Fig. 3B)**.

**Figure 3:**
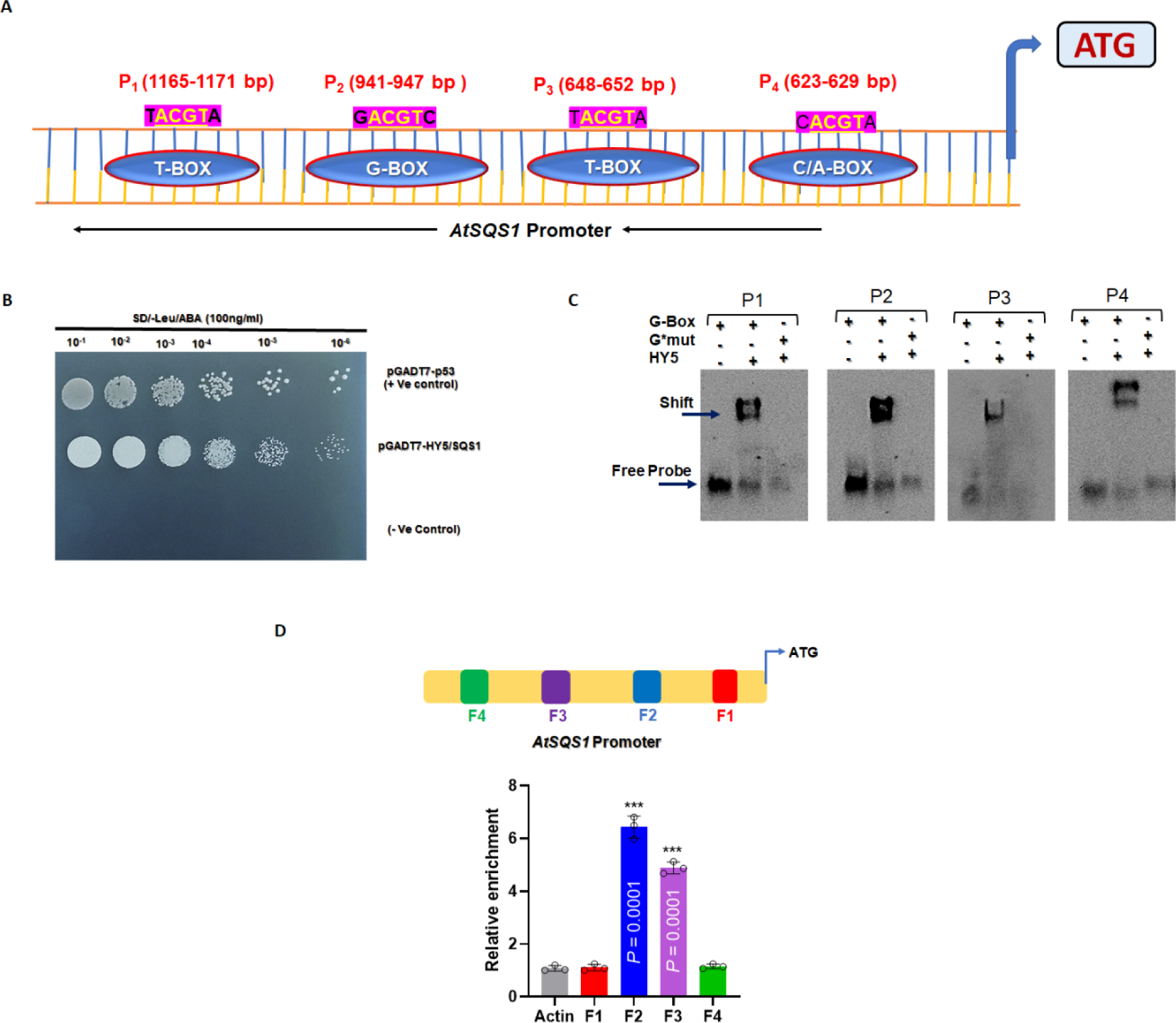
AtHY5 physically binds to the *AtSQS1* promoter. **(A)** Schematic representation of *AtSQS1* promoter with Light-responsive elements (LREs = G-BOX) located at four positions (P_1_) to (P_4_) in a region 2 kb upstream from the translational start site. **(B)** Yeast one-hybrid assay showing the interaction between HY5 and *AtSQS1* promoter. **(C)** EMSA for the binding of 6x-His-AtHY5 protein with a P_1_, P_2_, P_3_, and P_4_ digoxigenin-labeled probe (G-box) present in the *AtSQS1* promoter. **(D)** Promoter region showing primer location for each putative binding site used for Chip assay. Graph showing relative enrichment of the fragment. *Actin* was used as a negative control. The statistical analysis was performed using two-tailed Student’s t-tests. The data are plotted as means ± s.d. The error bars represent standard deviations. The asterisks indicate significant differences: *P< 0.001.

To further confirm the interaction between AtHY5 and the *AtSQS1* promoter, four probes were designed based on specific motifs within the promoter region (P1: 1165–1171 bp, P2: 941–947 bp, P3: 648–652 bp, and P4: 623–629 bp) and used in Electrophoretic Mobility Shift Assays (EMSA) **(Supplemental Fig. S4A)**. The assays were conducted using purified recombinant 6X-His-tagged AtHY5 protein, which exhibited strong binding to all four probes **(Fig. 3C)**. No binding was observed with mutated versions of these probes, indicating the specificity of the interaction. Competitive EMSA experiments were performed using both labeled and unlabeled (cold) probes to further verify binding specificity. When the labeled probes were incubated with increasing concentrations (10X to 50X) of the unlabeled probes, a reduction in the intensity of the shifted band was observed **(Supplemental Fig. S4C)**, confirming specific binding between AtHY5 and the *cis*-elements in the *AtSQS1* promoter. To validate this interaction *in-vivo*, chromatin immunoprecipitation followed by quantitative PCR (ChIP-qPCR) was conducted. AtHY5OX line revealed a significant enrichment of DNA fragments having the F2 and F3 regions, whereas no significant enrichment was observed for fragments containing the other two binding sites (F1 and F4) **(Fig. 3D).** These results provide evidence that AtHY5 directly binds to specific cis-regulatory elements found on the *AtSQS1* promoter, reinforcing its role as a negative regulator.

### Light-responsiveness of the *AtSQS1* promoter is dependent on AtHY5

The light responsiveness of the *AtSQS1* promoter was analyzed using histochemical GUS staining in promoter-reporter lines (*ProAtSQS1::*GUS\**GFP/*WT*, ProAtSQS1::*GUS\**GFP/hy5-215*, and *ProAtSQS1::GUS\**GFP/HY5OX) developed in WT*, hy5-215* mutant, and HY5OX backgrounds (**Supplemental Fig. S5A**). In WT and HY5OX backgrounds, GUS staining intensity decreased upon light exposure, whereas in the *ProAtSQS1::*GUS*GFP*/hy5-215* lines, GUS activity was not affected by light or dark conditions **(Fig. 4A)**. Similarly, GFP fluorescence in *ProAtSQS1::*GUS*GFP*/hy5-215* seedlings showed no change under light or dark conditions **(Fig. 4B)**. These findings suggest that *AtSQS1* is negatively regulated by light through HY5, as indicated by reduced GUS staining and GFP fluorescence in WT and HY5OX backgrounds under light exposure. Expression analysis showed that *AtCHS* (used as a positive control) exhibited a substantial increase under light exposure, while *AtSQS1* expression in *ProAtSQS1/*WT transgenic seedlings significantly decreased (**Fig. 4, C and D**). Moreover, dark-grown ProAtSQS1/WT and ProAtSQS1/HY5OX seedlings showed reduced GUS expression when shifted to light, whereas ProAtSQS1/*hy5-215* seedlings displayed no significant change in GUS expression under dark/ light conditions **(Fig. 4, E and F).** Control experiments with EV (EV used as positive control) and *ProAtSQS1*/*hy5-215* seedlings showed no change in GUS staining between light and dark conditions. In contrast, ProAtSQS1/WT and ProAtSQS1/HY5OX seedlings demonstrated a notable reduction in GUS activity upon light exposure compared to dark conditions **(Supplemental Fig. S5B).**

**Figure 4:**
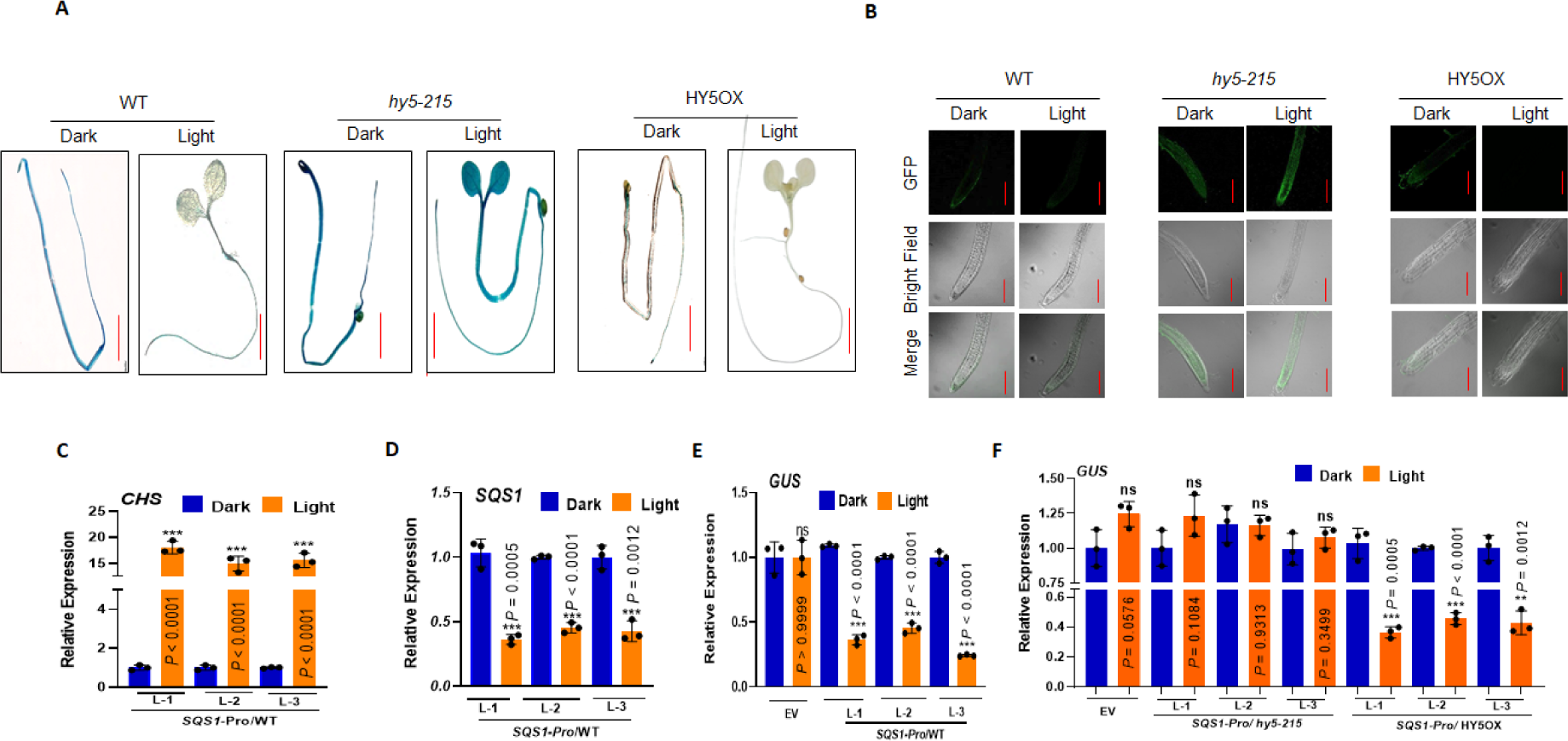
Light-responsiveness of the *AtSQS1* promoter in WT, *hy5-215*, and HY5OX backgrounds. **(A)** Histochemical staining showing GUS activity in five-day-old Arabidopsis seedlings transformed with EV, *ProAtSQS1::GUS*GFP/*WT, *ProAtSQS1::GUS*GFP/hy5-215* and *ProAtSQS1::GUS*GFP/*HY5OX. Scale bars, 1,000 µm. The experiment was repeated with three **(B)** Confocal images of 5-day-old seedlings grown in the presence of light and dark showing green fluorescence of GFP of *ProAtSQS1::GUS*GFP/*WT, *ProAtSQS1::GUS*GFP/hy5-215* and *ProAtSQS1::GUS*GFP/*HY5OX. Scale bars, 50 µM. Confocal images are representative of three independent experiments, n = 10 seedlings. (Scale bars, 50 μm). **(C and D)** RT-qPCR expression analysis of *GUS* reporter gene and levels of endogenous *AtSQS1* gene in seedlings of *ProAtSQS1::GUS*GFP/*WT, *ProAtSQS1::GUS*GFP/hy5-215* and *ProAtSQS1::GUS*GFP/*HY5OX grown on half-strength MS medium for 5 days in dark and light-grown seedlings. *Tubulin* was used as the endogenous control to normalize the relative expression levels. The statistical analysis was performed using two-tailed unpaired Student’s *t*-tests. Error bars represent the ± SD of means (*n* = 3). ***(P < 0.0001), **(P < 0.001), * (P < 0.05), ns (P > 0.05).

The responsiveness of the *AtSQS1* promoter was further validated through GUS histochemical staining in transgenic seedlings, demonstrating tissue-specific expression, particularly in the inflorescence and siliques **(Supplemental Fig. S6A)**. The staining results showed comparable expression levels in *ProAtSQS1::GUS*\*GFP*/hy5-215* and the EV control, indicating that the absence of functional AtHY5 in *ProAtSQS1::GUS*\*GFP/*hy5-215* plants led to a lack of negative regulation by light, resulting in increased GUS staining in the inflorescence and siliques. In contrast, *ProAtSQS1::GUS*\*GFP/WT and *ProAtSQS1::GUS*\*GFP/HY5OX mature plants exhibited minimal or negligible GUS staining, reflecting light-mediated negative regulation by functional light-responsive components **(Supplemental Fig. S6A)**. Additionally, GFP fluorescence was analyzed in the root tips of promoter-reporter lines seedlings harboring the *AtSQS1* promoter construct in WT, *hy5-215*, and HY5OX backgrounds under light and dark conditions **(Fig. 4B)**. In light conditions, GFP fluorescence was absent in the root tips of *ProAtSQS1::GUS*\*GFP/WT and *ProAtSQS1::GUS*\*GFP/HY5OX seedlings but was detected in *ProAtSQS1::GUS*\*GFP*/hy5-215* lines. A similar GFP expression pattern was observed in the root elongation zone of these seedlings **(Supplemental Fig. S6B)**.

GUS histochemical analysis was performed at various time intervals following light exposure to study GUS accumulation in transgenic seedlings. In *ProAtSQS1::GUS*\*GFP/WT and *ProAtSQS1::GUS*\*GFP/HY5OX lines, GUS accumulation progressively decreased from 30 minutes to 24 hours post-exposure. In contrast, EV (EV/WT; EV/AtHY5OX) plants showed no significant change in GUS levels during the transition from dark to light **(Supplemental Fig. S7A)**. Notably, *ProAtSQS1::GUS*\**GFP*/*hy5-215* transgenic seedlings displayed unchanged GUS accumulation across the light exposure intervals, indicating a lack of response due to the absence of functional HY5. These findings were supported by the relative expression analysis of the GUS reporter gene, which was similar to histochemical staining results **(Supplemental Fig. S7B)**.

### *AtSQS1* overexpression enhances root growth and metabolite accumulation

The analysis of root length phenotypes in various genetic backgrounds of *AtSQS1*-overexpressing plants revealed significant insights into the regulatory role of AtHY5 in plant development and metabolite content. In 10-day-old seedlings grown on half-strength MS medium, SQS1OX/WT, SQS1OX/*sqs1*, SQS1OX/*hy5-215*, and SQS1OX/HY5OX lines exhibited enhanced root growth compared to their corresponding control backgrounds (WT, *sqs1*, *hy5-215*, and HY5OX, respectively) **(Fig. 5, A and B)**. Quantitative expression analysis confirmed that *AtSQS1* expression was markedly elevated in all SQS1OX lines compared to their controls **(Fig. 5C)**. Furthermore, total terpenoid quantification revealed that SQS1OX/*hy5-215* plants had the highest terpenoid content, whereas SQS1OX/*sqs1* lines displayed the lowest content, however, this was higher compared to the wild type. **(Supplemental Fig. S8)**. These findings indicate that HY5 acts as a negative regulator of *AtSQS1* expression and terpenoid biosynthesis, while *AtSQS1* overexpression promotes terpenoid accumulation.

**Figure 5:**
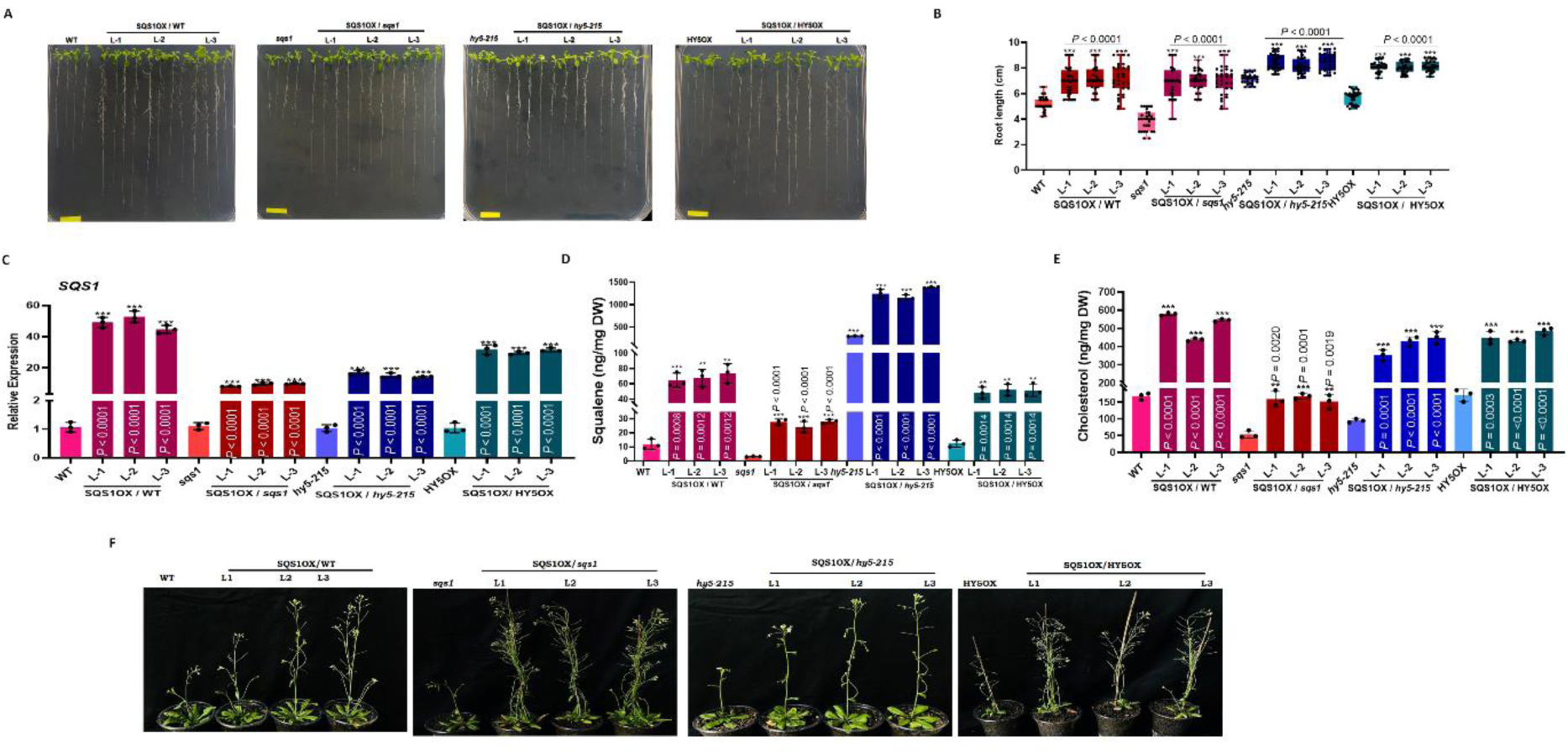
The overexpression of AtSQS1 in Arabidopsis leads to the accumulation of metabolites and promotes growth. **(A)** Representative image of 10-day-old overexpression lines in different backgrounds of light-modulated lines SQS1OX/WT, SQS1OX/*sqs1*, SQS1OX/*hy5-215*, and SQS1OX/HY5OX seedlings grown on half-strength MS medium. Scale bars, 1 cm. **(B)** RT-qPCR of SQS1 in the WT, *hy5-215,* SQS1OX/WT, SQS1OX/*sqs1*, SQS1OX/*hy5-215*, and SQS1OX/HY5OX lines. **(C)** Graphical representations of the WT, *hy5-215,* SQS1OX/WT, SQS1OX/*sqs1*, SQS1OX/*hy5-215*, and SQS1OX/HY5OX seedlings grown on half-strength MS medium. *n* = 30; independent seedlings, small open circles. **(D and E)** Quantification of squalene and cholesterol content levels in the WT, *hy5-215,* SQS1OX/WT, SQS1OX/*sqs1*, SQS1OX/*hy5-215*, and SQS1OX/HY5OX seedlings **(F)** Representative images of 30-day-old light-modulated lines SQS1OX/WT, SQS1OX/*sqs1*, SQS1OX/*hy5-215*, and SQS1OX/HY5OX plants (L1, L2, L3). Plants were grown under a 16-h-light/8-h-dark photoperiod. The statistical analysis was performed using two-tailed Student’s *t*-tests (n= 30). The data are plotted as means ±SD between the treatment and the control. The error bars represent standard deviations. The asterisks indicate significant differences: *P< 0.01; **P< 0.001; ***P< 0.0001.

Quantification of squalene and cholesterol content was performed in SQS1OX/WT, SQS1OX/*sqs1*, SQS1OX/*hy5-215*, and SQS1OX/HY5OX lines, along with their respective control backgrounds. The analysis of squalene levels demonstrated a pattern similar to total terpenoid content, showing the highest accumulation in SQS1OX/*hy5-215* and the lowest in SQS1OX/*sqs1* plants **(Fig. 5D)**. Squalene content in SQS1OX/WT and SQS1OX/HY5OX lines were comparable. These results indicate that AtSQS1 positively regulates squalene biosynthesis, while HY5 functions as a negative regulator. Cholesterol levels were enhanced in all SQS1OX lines relative to their respective controls **(Fig. 5E)**.

To assess the effect of *AtSQS1* overexpression on growth parameters such as bolting time, rosette diameter, and plant height, these traits were analyzed in SQS1OX/WT, SQS1OX/*sqs1*, SQS1OX/*hy5-215*, and SQS1OX/HY5OX lines at the mature stage (30 days post-sowing). Quantitative expression analysis confirmed significantly higher levels of AtSQS1 in all SQS1OX lines compared to their corresponding control backgrounds **(Supplemental Fig. S9, A-B and Supplemental Fig. S10, A-B)**. As a result, early bolting was observed in SQS1OX/WT, SQS1OX/*sqs1*, SQS1OX/*hy5-215*, and SQS1OX/HY5OX plants, as it occurred 5-6 days earlier than in their respective control backgrounds. Phenotypic measurements demonstrated that rosette diameter and plant height were substantially greater in the SQS1OX lines compared to their controls **(Supplemental Fig. S9, C-E and Supplemental Fig. S10, C-E)**. Additionally, the analysis of phytosterol content revealed significantly elevated levels of stigmasterol, campesterol, cycloartenol, and β-sitosterol in SQS1OX/WT, SQS1OX/*sqs1*, SQS1OX/*hy5-215*, and SQS1OX/HY5OX plants relative to their controls **(Supplemental Fig. S11, A-D)**. These findings indicate that *AtSQS1* overexpression not only promotes early bolting and enhances growth but also positively influences phytosterol biosynthesis. Out of four phytosterols analysed in our study, the content of β-sitosterol was highest in all the above-mentioned lines (Michael et al. 2024).

### Mutation in *AtSQS1* in WT and *hy5-215* backgrounds affects plant development

Overexpression of *AtSQS1* in different genetic backgrounds led to significant enhancement in plant growth and development. To validate these results, CRISPR/Cas9-based genome-edited knockout mutants of *AtSQS1* in WT and *hy5-215* backgrounds *(SQS1^CR^/*WT and *SQS1^CR^/hy5-215)* were developed, with nucleotide insertions and deletions **(Supplemental Fig. S12A)**. The height of 30-day-old plants of *SQS1^CR^/*WT and *SQS1^CR^/hy5-215* lines was very low as compared to their respective controls, i.e., WT and *hy5-215* plants (**Fig. 6A**). This phenotype was further supported by expression analysis. Expression analyses revealed significantly reduced *AtSQS1* transcript levels in *SQS1^CR^/*WT and *SQS1^CR^/hy5-215* mutant plants compared to their respective control backgrounds **(Fig. 6B)**. The *SQS1^CR^/*WT and *SQS1^CR^/hy5-215* mutant lines exhibited reduced plant height, smaller rosette diameter, and delayed bolting compared to their respective control plants **(Fig. 6A, Supplemental Fig. S15, A-C)**. Interestingly, the characteristic elongated hypocotyl phenotype of the *hy5-215* mutant was absent in *SQS1^CR^/hy5-215* plants. *SQS1^CR^/hy5-215* plants displayed a distinctive phenotype as compared to WT i.e., their hypocotyls were shorter and stunted than the control (**Supplemental Fig. S14)**. Western blot analysis was conducted to validate this altered phenotype of *SQS1^CR^/hy5-215* plants by confirming the absence of the HY5 protein in the *SQS1^CR^/hy5-215* plants **(Fig. 6C and Supplemental Fig. S14)**. Metabolite profiling showed a significant reduction in squalene and cholesterol content in these mutants relative to their control **(Fig. 6, D and E)**. Notably, squalene levels were significantly reduced in *SQS1^CR^/hy5-215* compared to *hy5-215* **(Supplemental Fig. S12B)**. Further investigation into the expression of key terpenoid biosynthetic genes revealed downregulation of *AtSQE3*, *AtDWF5*, and *AtCAS1* in both *SQS1^CR^/*WT and *SQS1^CR^/hy5-215* mutants compared to their controls **(Supplemental Fig. S13, A and B)**. The mutation in *AtSQS1* in both WT and *hy5-215* backgrounds significantly impacted plant growth and morphology.

**Figure 6.**
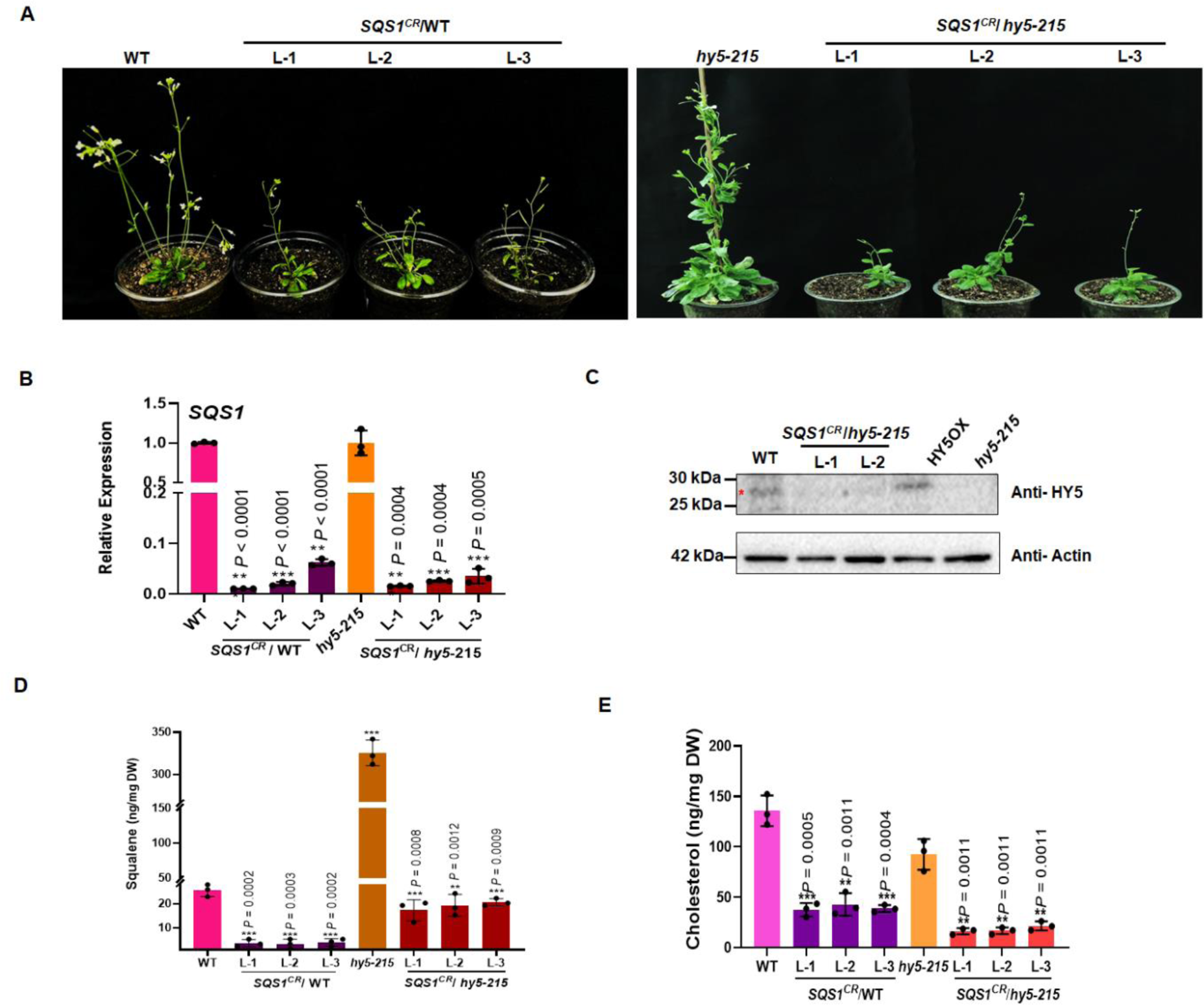
*AtSQS1*CRISPR/Cas9 edited lines in *hy5-215* mutants and WT plants show altered phenotype and modulated metabolites contents. (A) Representative image of 30-day-old *hy5-215*, *SQS1^CR^*/*hy5-215,* and WT, *SQS1^CR^*/WT plants. **(B)**Quantification of SQS1 in *SQS1^CR^*/*hy5-215* and *SQS1^CR^*/ WT (the small open circles represent the individual values). This experiment was repeated three times independently, with similar results. **(C)**Western blot analysis of HY5 in five-day-old seedlings of WT, HY5OX, *hy5-215*, and *SQS1^CR^/hy5-215* edited plants. Actin was used as the loading control. The experiment was repeated three times independently with similar results. **(D and E)** Quantification of Squalene and cholesterol contents in *hy5-215*, *SQS1^CR^*/*hy5-215,* and WT, *SQS1^CR^*/WT through GC-MS. The statistical analysis was performed using two-tailed Student’s t-tests. The data are plotted as means ± s.d. The error bars represent standard deviations. The asterisks indicate significant differences: *P< 0.001.

### Exogenous squalene restores *AtSQS1^CR^* mutant phenotype in different background

To investigate the potential of exogenous squalene in restoring *AtSQS1* function in *SQS1^CR^/*WT and *SQS1^CR^/hy5-215* mutants, root length measurements were performed on seedlings grown on media supplemented with various squalene concentrations (100–1000 ng/L). The root lengths of *AtSQS1^CR^/*WT and *AtSQS1^CR^/hy5-215* plants were significantly shorter than those of their respective control backgrounds (WT and *hy5-215*) **(Fig. 7, A-C)**. However, supplementation of exogenous squalene led to a significant increase in root length in the mutant seedlings. The root growth enhanced significantly when 300 ng/l squalene was administered exogenously whereas concentrations exceeding 500 ng/l adversely affected root length and overall growth. On the other hand, when exogenous squalene was supplemented with dark-grown seedlings of the *AtSQS1^CR^*/WT, no significant modulation was observed in plant growth **(Supplemental Fig. S16)**. These above results clearly suggest the role of light in plant growth and development via squalene and other phytosterols.

**Figure 7:**
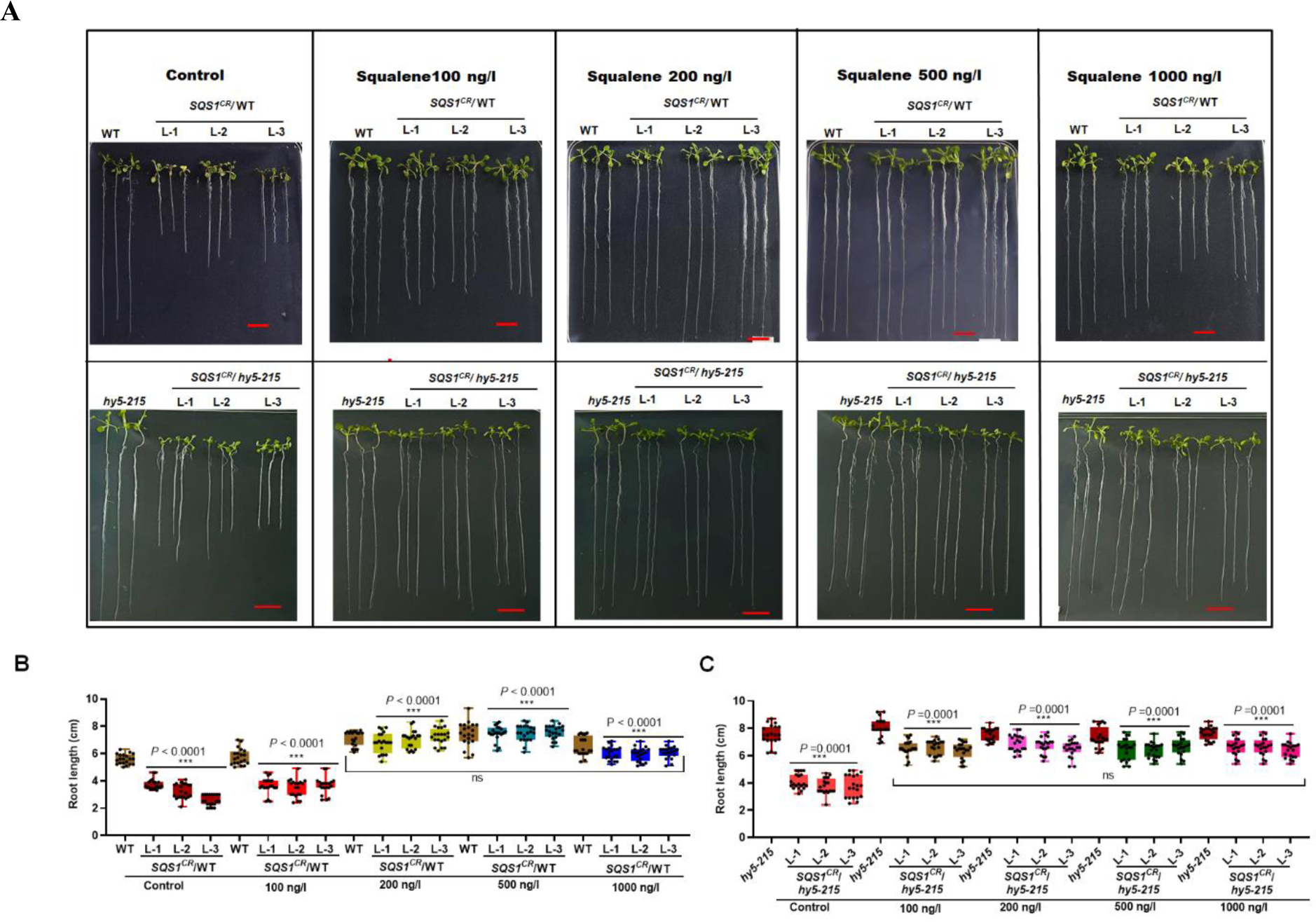
Exogenous squalene restores seedling root growth in CRISPR/Cas9-based edited mutant plants of *SQS1^CR^/hy5-215* and *SQS1^CR^*/WT. **(A)** Representative image of ten-day-old seedlings of WT, *hy5*-215, *SQS1^CR^*/WT and *AtSQS1^CR^*/*hy5-215* lines grown on half-strength MS medium supplemented with water (control) and 100-1000 ng/l squalene, Scale bar, 1 cm. **(B and C)** Root lengths of ten-day-old seedlings of WT, *hy5*-215, *SQS1^CR^*/WT and *AtSQS1^CR^*/*hy5-215* lines grown on half-strength MS medium supplemented with water (control) and 100 to 1000 µM squalene, (n= 20 independent seedlings, indicated by the small open circles). The experiment was repeated three times independently, with similar results. The statistical analysis was performed using two-tailed Student’s *t*-tests (n= 20). The data are plotted as means ±SD between the treatment and the control. The error bars represent standard deviations. The asterisks indicate significant differences: \**P*< 0.01; \*\**P* < 0.001; \*\*\**P* < 0.0001.

## Discussion

Previous studies have extensively documented the role of light in plant growth, development, and flavonoid biosynthesis (Yadav et al. 2020; Bhatia et al. 2021; Lui et al. 2023; Lee et al. 2024; Singh et al. 2024). However, the regulation of terpenoid biosynthesis by light still remains less understood (Shi et al. 2022; Hernandez et al. 2023; Wei et al. 2023; Michael et al. 2024). Research in *Arabidopsis thaliana* has shown that light positively influences the transcript levels of genes in the MEP pathway, which is crucial for the synthesis of photosynthetic pigments during early seedling development (Banerjee and Sharkey 2014; Chen et al. 2020). This regulatory mechanism ensures an increased supply of photosynthetic pigments derived from the MEP pathway, supporting plant growth and photosynthetic efficiency. Despite these insights, there remains a gap in understanding how light-responsive signal transduction pathways and specific transcription factors modulate the terpenoid biosynthesis under varying environmental conditions (Michael et al. 2020; Michael et al. 2024). This study addresses this gap by investigating the MVA pathway and its sterol biosynthesis branch, emphasizing their regulation by light. Our findings highlight that light significantly impacts the transcriptional regulation of terpenoid biosynthesis, particularly influencing the synthesis of squalene and phytosterols in *Arabidopsis thaliana*. We further identified that *AtSQS1* expression is regulated by light, with the transcription factor HY5 playing a key mediatory role.

To elucidate the role of light in regulating the gene expression of the squalene biosynthesis pathway and its downstream sterol branch, an *in-silico* promoter analysis was performed **(Supplemental Fig. S4A)**. This analysis revealed the presence of multiple putative light-responsive elements (LREs) in the *AtSQS1* promoter region. In dark-grown WT seedlings, exposure to light led to a substantial decrease in both *AtSQS1* transcript levels and squalene content, indicating light-mediated repression. Time-course analysis further demonstrated a progressive decline in *AtSQS1* expression with extended light exposure **(Fig. 1)**. In dark condition, *hy5-215 and* AtHY5OX plants both have long hypocotyls as compared to the WT. The possible reason for this phenotype is that AtHY5OX plants were developed in the *hy5-215* mutant background (AtHY5OX/*hy5-215;* Bhatia et al. 2018). In *hy5-215* mutant seedlings, *AtSQS1* expression and squalene content were significantly enhanced compared to WT, suggesting that the absence of functional HY5 relieves repression. Conversely, AtHY5OX seedlings exhibited significantly lower *AtSQS1* transcript levels and squalene content, indicating that increased HY5 levels negatively impact squalene biosynthesis **(Fig. 2)**. Exposing the dark-grown AtHY5OX seedlings to light conditions resulted in a significant reduction in *AtSQS1* expression, as well as squalene and overall terpenoid content. Interestingly, the cholesterol content was high in *hy5-215* plants as compared to the WT. Though there are no reports available about HY5 regulating cholesterol biosynthesis, we speculate that increased cholesterol content could be due to increased squalene in these plants, as cholesterol is also a downstream metabolite of the same pathway **(Fig. 2G).** According to a previous report, UV-B light plays an important role in the biosynthesis and regulation of vitamin D3, a sitosterol which is produced by the photolytic conversion of cholesterol biosynthetic intermediate provitamin D3 in *Arabidopsis thaliana* (Silvestro et al. 2018). This report suggests the role of light components in cholesterol biosynthesis, but detailed experimental validation and research are needed to confirm these findings. These findings highlight HY5 as a key negative regulator of *AtSQS1* expression, demonstrating that light-dependent modulation of squalene biosynthesis is mediated through HY5.

To validate the regulation of *AtSQS1* gene expression by HY5, the interaction between the *AtSQS1* promoter and HY5 was examined using EMSA and Y1H assays. The EMSA results revealed strong binding and band shifts when the 6X-His-AtHY5 protein was incubated with four probes (P1, P2, P3, and P4), which were designed from the upstream genomic region (∼1.9 kb upstream of the translational start site) of the *AtSQS1* promoter, specifically targeting the G-box motif **(Fig. 3C)**. This observation aligns with previous studies demonstrating that HY5 can transcriptionally regulate other genes, such as *AtTPS03*, by binding to their promoters (Michael et al. 2020). Further validation was conducted through ChIP assays, which confirmed that HY5 binds to specific light-responsive elements within the *AtSQS1* promoter. Histochemical analysis of *AtSQS1Pro::GUS/*HY5OX transgenic seedlings grown in darkness for 5 days, followed by exposure to white light, revealed a progressive decrease in GUS activity with increasing durations of light exposure **(Fig. 4F and Supplemental Fig. S5B)**. This reduction in GUS expression was observed consistently across various plant tissues, including seedlings, inflorescences, and siliques **(Supplemental Fig. S6A)**. Notably, transgenic seedlings on the HY5OX background exhibited significantly reduced GUS expression compared to those on the WT background **(Fig. 4F and Supplemental Fig. S7B)**. These findings suggest that HY5 negatively regulates *AtSQS1* promoter activity, reinforcing the role of HY5 as a transcriptional repressor of squalene biosynthesis in response to light. These combined findings provide evidence for the direct physical interaction of HY5 with the G-box *cis*-regulatory elements in the *AtSQS1* promoter, indicating that HY5 plays a critical role in modulating *AtSQS1* transcription in response to light signals.

Plant sterols and their derivatives play crucial roles in regulating key physiological processes, including seed germination, flowering time, senescence, and seed yield. These compounds are essential for maintaining the permeability, fluidity, and integrity of cellular membranes (Simons et al. 2011; Rogowska et al. 2020). We analysed the physiological parameters of the developed *AtSQS1* overexpression lines and mutants in different genetic backgrounds. In transgenic seedlings overexpressing *AtSQS1*, root length was significantly increased compared to their respective controls across all backgrounds, including WT and HY5OX **(Fig. 5, A and B)**. In addition, bolting time, rosette diameter, and plant height in *AtSQS1OX* plants were altered compared to their respective controls, with these plants maturing earlier and completing their life cycle in a shorter period. Metabolite analysis revealed significant increases in terpenoids (squalene) and plant phytosterols (cholesterol, stigmasterol, campesterol, β-sitosterol, and cycloartenol) in the overexpression lines (SQS1OX*/sqs1*, SQS1OX*/*WT, SQS1OX*/hy5-215*, and SQS1OX/HY5OX) compared to their respective control backgrounds **(Supplemental Fig. S11)**. These findings align with previous studies, such as the *ssr1* mutant in *Arabidopsis thaliana*, which exhibited reduced production of sitosterol, stigmasterol, and campesterol, leading to abnormal stigma development and partial sterility (Vriese et al. 2021). Additionally, plant sterols, such as brassinolide, are critical for plant structure, and disruption of the SMT2 gene in the sterol biosynthesis pathway results in a pronounced dwarf phenotype, a hallmark of BR-deficient mutants. This phenotype has been documented in several species, including cherry tomato (Rahim et al. 2018), *Arabidopsis* (Klahre et al. 1998), rice (Hong et al. 2002), and pea (Nomura et al. 2004). Taken together, these results support our data, suggesting that enhanced sterol content in AtSQS1OX plants contributes to improved growth parameters and overall plant development.

We also generated *AtSQS1* mutants using a CRISPR/Cas9-based genome-editing approach in both WT and *hy5-215* backgrounds. In these mutants (*SQS^CR^/*WT and *SQS^CR^/hy5-215*), the expression of *AtSQS1*, squalene content, and cholesterol content were significantly reduced in both seedlings and mature plants **(Fig. 6, B-E)**. Furthermore, the expression levels of genes involved in squalene biosynthesis, including *AtSQE3*, *AtCAS1*, and *AtDWF5*, were also downregulated in the mutants **(Supplemental Fig. S13)**. The mutants exhibited reduced plant height and rosette diameter compared to their respective control backgrounds, reflecting altered growth and developmental phenotypes. These findings are consistent with previous studies, where mutations in genes of the MVA pathway, such as *DWF5* and *HMGS*, led to delayed seed germination and morphological alterations. In *Arabidopsis thaliana*, the *dwf5* and *dwf7* mutants, which lack sterols, exhibit significant inhibition of seed germination and noticeable changes in seed morphology (Yu et al. 2021). Additionally, *A. thaliana* seedlings overexpressing *HMGS* have been shown to germinate faster and accumulate higher sterol levels compared to wild-type plants. These results highlight the importance of sterols in plant growth and development and further emphasize the role of *AtSQS1* and its downstream biosynthetic pathways in regulating key physiological processes. Our results also suggest that *SQS1^CR^/hy5-215 mutants* had short and stunted hypocotyls; as these plants lack both AtSQS1 and AtHY5 protein, they have a phenotype very similar to wild type **(Fig. 6C)**. The absence of the HY5 protein in the *SQS1^CR^/hy5-215* plants was further confirmed by western blot analysis (**Supplemental Fig. S14)**

To assess whether the application of exogenous metabolites could restore the mutant phenotype, seedlings of *AtSQS1^CR^/hy5-215* plants were grown on media supplemented with squalene at varying concentrations. The *AtSQS1^CR^/hy5-215* mutant lines, which displayed significantly shorter root lengths compared to their respective controls, demonstrated restoration of root length to levels comparable to non-mutant plants upon squalene treatment **(Fig. 7, A-C)**. This phenotype recovery aligns with prior findings in which the dwarf phenotype of the *Pisum sativum smt2* mutant, characterized by brassinosteroid (BR) deficiency, was reversed through the exogenous application of brassinolide. Such results underscore the potential of targeted metabolite supplementation to mitigate growth defects associated with disruptions in sterol and terpenoid biosynthetic pathways. Our results further indicate the role of squalene in plant growth and development through light-dependent phenomenon as exogenous squalene when supplemented to dark-grown seedlings of the *AtSQS1^CR^*/WT leads to no signification alteration in root phenotype **(Supplemental Fig. S16)**.

Based on our findings, we propose a regulatory model **(Fig. 8)** illustrating how HY5 directly interacts with G-box *cis*-regulatory elements in the *AtSQS1* promoter, highlighting its pivotal role in modulating *AtSQS1* transcription in response to light signal. In the dark, the absence of HY5, which acts as a negative regulator of squalene biosynthesis, allows the increased *AtSQS1* expression, resulting in higher squalene accumulation. Conversely, under light conditions, HY5 binds to the *AtSQS1* promoter and represses its transcription, thereby leading to a reduction in squalene levels. This model underscores the light-dependent regulatory mechanism by which HY5 modulates squalene biosynthesis, aligning plant metabolic output with environmental light conditions to optimize growth and development.

**Figure 8:**
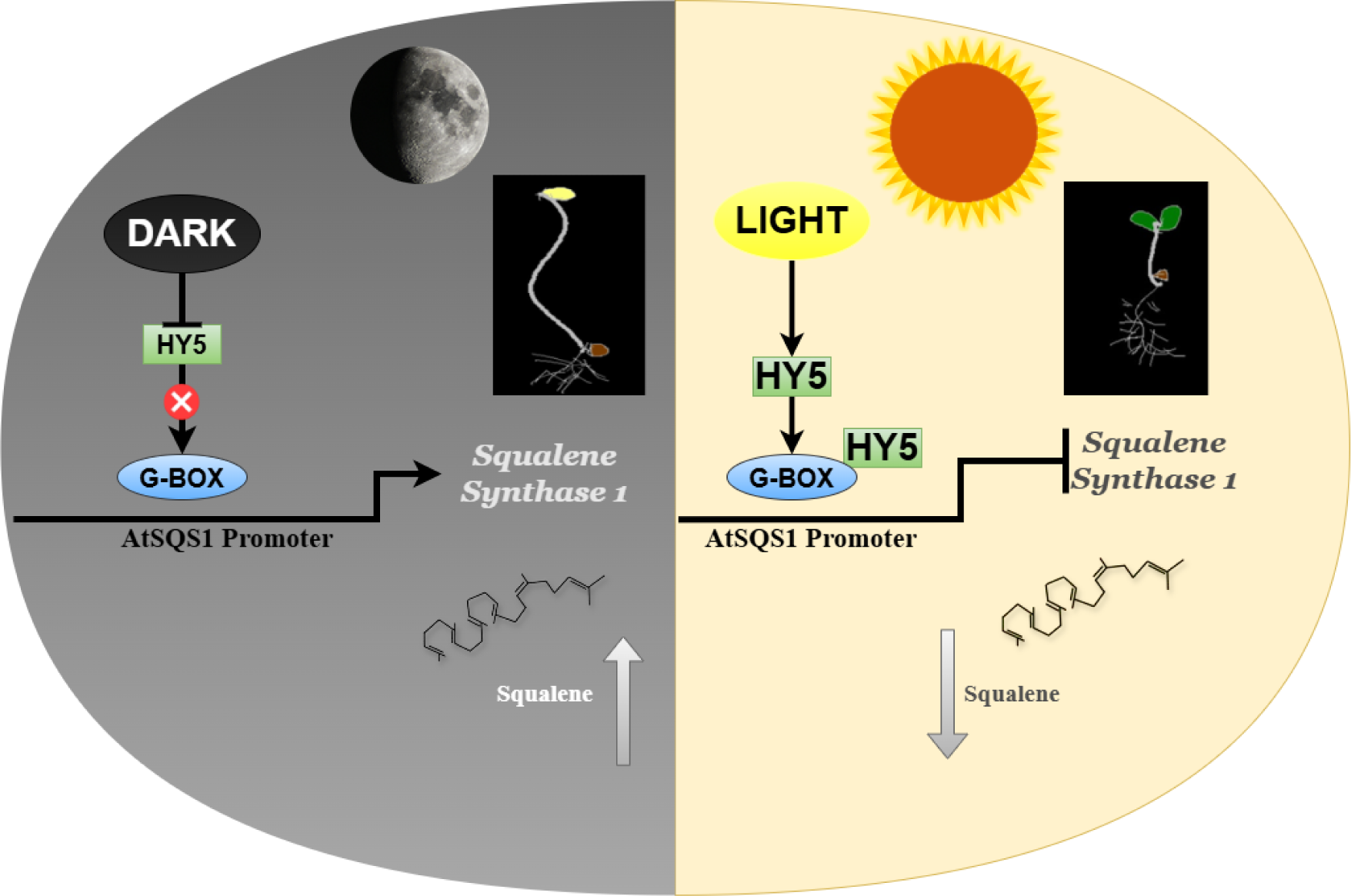
Proposed model for light-mediated regulation of squalene biosynthesis in *A. thaliana* regulated by HY5. In the dark, nuclear-localized proteasomal complex targets HY5 for degradation and prevents photomorphogenesis. HY5 does not bind on the SQS1 promoter and thus enhances squalene biosynthesis. In the light, active HY5 is present in the nucleus and binds to the promoter of SQS1, leading to the inhibition of squalene production. Light negatively regulates *Squalene synthase 1*, and dark conditions favor squalene production.

## Material methods

### Plant materials and growth conditions

*Arabidopsis thaliana* accession Columbia-0 (Col-0, CS60000), the wild-type (WT henceforward) employed in all experiments within this study. The mutants *sqs1* (SALK-077057C)*, hy5-215,* and used were previously described (Oyama et al. 1997; Johnson et al. 2014). Transgenic/mutant (SQS1OX/WT, SQS1OX/*sqs1,* SQS1OX/*hy5-215*, SQS1OX/HY5OX, *SQS1^CR^*/WT, *SQS1^CR^/hy5-215*) plants were developed and utilized in this study. HY5OX lines were developed earlier by our group and used in this study for analysis (Bhatia et al. 2018). The seeds were surface-sterilized and plated on half-strength Murashige and Skoog (MS) medium (Hi-media) supplemented with 1.5 % sucrose. After stratification for 3 days at 4 °C in the dark, the plates were transferred to a growth chamber (Percival) maintained under a long-day photoperiod cycle (16-h light, 8-h dark) and constant temperature (22°C). The plates were covered in 2-3 layers of aluminum foil for complete darkness treatment and harvested in dim green light.

### *In silico* analysis of promoter

For cis-regulatory elements, an *in silico* analysis of the promoter was conducted. The promoter sequences of AtSQS1, approximately 1.8 kb upstream of the translation start site, were obtained from the AGRIS database (Palaniswamy et al. 2006; Yilmaz et al. 2010). and PlantPAN3.0 (Chow et al. 2019). LREs were identified both manually and through the use of PlantPAN3.0 (https://plantpan.itps.ncku.edu.tw/plantpan3).

### Yeast one-hybrid analysis

For Y1H assay, up to 1.9 kb upstream promoter region of *AtSQS1* consisting of all LREs, specifically core G-BOX elements, was targeted. The above-mentioned promoter was amplified from genomic DNA and cloned into pAbAi plasmid to construct bait construct: *ProAtSQS1-pABAi*. HY5 sequences were cloned into pGADT7-AD plasmid to construct prey vectors with translational fusions with activation domain (pGADT7AD-AtHY5) sequence-specific primers. Further, prey constructs AtHY5-AD were co-transformed into Y1H gold*-ProAtSQS1-pAbAi* bait strain. The positive interactions between the bait *ProAtSQS1*-*pAbAi* and the prey pGADT7AD-*AtHY5* were confirmed by spotting these strains on SD/-leu AbA (200 ng/ml) agar medium at *cfu* 1X10^3^ cells/ml. The interaction between *Prop53-pAbAi* and p53-AD was crucial as a positive control in the study.

### Electrophoretic mobility shift assay

EMSA analysis and labelling of probes with digoxigenin were done using the 2nd generation DIG Gel Shift EMSA kit (Roche, Sigma Aldrich; USA) as per the manufacturer’s instructions. Labelled probes were incubated for 30 min at 20^0^C in binding buffer [100 mM HEPES (pH 7.6), 5 mM EDTA, 50 mM (NH_4_)_2_SO_4_, 5 mM DTT, Tween 20, 1% (w/v), 150 mM KCl] with or without recombinant protein. In a brief reaction, 20 µl mixture containing binding buffer, 1 µg/μl Poly (dI-dC), 15-fmol/μl DIG-labelled probes with varying amounts of the unlabelled probe as a competitor and purified protein (∼2 µg). The binding reaction was resolved on a 6 % polyacrylamide gel in 0.5X TBE buffer and was semi-dry blotted (Transblot, BIO-RAD, USA) onto a positively charged nylon membrane (BrightStar, Invitrogen, USA). The membrane was UV cross-linked and was developed as per the manufacturer’s instructions. The developed blot was finally incubated with CSPD chemiluminescent solution and exposed to X-ray blue film (Retina, India). The positively charged nylon membrane was incubated with CSPD chemiluminescent solution, and images were captured using Image Lab version 5.2.1 build 11 (Bio-Rad Laboratories).

### ChIP analysis

Ten-day-old seedlings were used to perform chromatin immunoprecipitation (ChIP). Seedling tissues (∼2 g) were cross-linked with 50 mL of 1% formaldehyde in a vacuum for 20 minutes. To stop the cross-linking, 2.5 mL of 2 M glycine was added. Seedlings were rinsed with water, ground in liquid nitrogen, and resuspended in 20 mL of Extraction Buffer 1, containing 0.4 M sucrose, 10 mM Tris–HCl (pH 8), 10 mM MgCl_2_, 5 mM beta-mercaptoethanol, 0.1 mM PMSF, and a protease inhibitor (Sigma). The mixture was then filtered through a cell strainer (Corning). The filtrate was centrifuged at 4,000 rpm for 30 minutes at 4 °C. The pellet was resuspended in 1 mL of Extraction Buffer 2, which included 0.25 M sucrose, 10 mM Tris–HCl (pH 8), 10 mM MgCl_2_, 1% Triton X-100, 5 mM beta-mercaptoethanol, 0.1 mM PMSF, and a protease inhibitor, and then centrifuged at 14,000 rpm for 10 minutes. Next, the pellet was resuspended in 300 mL of Extraction Buffer 3, containing 1.7 M sucrose, 10 mM Tris–HCl (pH 8), 0.15% Triton X-100, 2 mM MgCl_2_, 5 mM beta-mercaptoethanol, 0.1 mM PMSF, and protease inhibitors. This solution was loaded on top of an equal volume of clean Extraction Buffer 3 and centrifuged at 14,000 rpm for 1 hour. The crude nuclear pellet was resuspended in nuclear lysis buffer (50 mM Tris–HCl, pH 8.0; 10 mM EDTA; 1% SDS; and complete protease inhibitor from Roche) and sonicated to achieve fragment sizes of 0.3 to 0.8 kb. After centrifugation, the insoluble pellet was discarded. The soluble chromatin was diluted 10-fold with ChIP dilution buffer (1.1% Triton X-100, 1.2 mM EDTA, 16.7 mM Tris–HCl, pH 8.0, and 167 mM NaCl). Following pre-clearing with protein-A Sepharose beads (Sigma-Aldrich), 40 µL of HA tag-specific monoclonal antibody (Roche) was added and incubated overnight at 4 °C. Immunocomplexes were extracted using 100 µL of 50% protein-A Sepharose beads for 1 hour at 48 °C. After washing, the immunocomplex was eluted twice with 250 µL of elution buffer (1% SDS and 0.1 M NaHCO_3_) and reverse cross-linked at 65 °C for 12 hours with 200 mM NaCl. After protein removal with proteinase K, DNA was purified using phenol-chloroform extraction and ethanol precipitation. The pellet was resuspended in 50 µL of 0.13 TE (10 mM Tris-EDTA, pH 7.5) with RNase A (0.1 mg/mL) for probe synthesis or PCR analysis.

### Construct preparation and development of overexpressing/mutant plants

For the editing of *AtSQS1* using the CRISPR-Cas9 system in *A. thaliana*, gRNA from coding region of AtSQS1 was designed using the CRISPR Arizona software (http://www.genome.arizona.edu/crispr). gRNA was cloned into the binary vector pHSE401 using the *Bsa*I restriction site (Sharma et al. 2020). This vector contains Cas9 endonuclease-encoding gene under dual CaMV35S promoter as well as genes encoding neomycin phosphotransferase and hygromycin phosphotransferase as selection markers. The construct was sequenced from both the orientations using plasmid-specific forward and vector reverse primers. For the overexpression of SQS1, the complete CDS was amplified using cDNA of Col-0. The amplicon was cloned into pTZ57R/T and then transferred into plant expression vector pCAMBIA1301 containing the CaMV35S promoter. For the histochemical assay, promoter lines were developed with the GUS reporter gene, using around 1.9-kb upstream sequence from the translation initiation codon. The amplicons were cloned into pCAMBIA1303 plant binary vector. The plant transformation vectors pCAMBIA1301/pCAMBIA1303 carrying At*SQS1* and pHSE401 with gRNA were used to transform the *Agrobacterium* strain GV3101. Arabidopsis thaliana (Col-0/ *hy5-215*/HY5OX/, *sqs1*) plants were used for transformation by using the floral dip method (Clough and Bent, 1998).

The transformants were screened on a half-strength MS agar plate containing the appropriate antibiotics. Putative mutated transformants were identified as hygromycin-resistant seedlings that produced green leaves and well-established roots in the selection medium, while in kanamycin, non-transformants displayed a typical chlorosis symptom. After 8-10 days, each viable seedling was transferred into individual pots filled with soilrite and saturated with nutrient media. Genomic DNA from the leaves was isolated using the CTAB method. Regions flanking the target site were amplified by PCR, and mutations were detected using the Takara Guide-it Mutation Detection Kit (Takara). PCR products from positive plants were cloned in the cloning vector pTZ57R/T and sequenced using M13F and M13R primers.

### Phenotypic analysis

The phenotypic parameters were carefully observed and consistently monitored for up to 2-3 generations. For studying the variation at the seedling level, 40-50 seeds of transgenic lines along with WT were surface-sterilized and germinated on a medium consisting of ½MS. Plates were kept at 4°C for 2 days, replaced in a growth chamber at a constant 22°C under 16 h light/8 h dark period, and grown vertically. After 10 days of seedling growth, their root length was recorded along with the roots of WT plants grown simultaneously under similar conditions. Mature plants (30 days and 35 days) of WT and transgenic lines were observed for plant height, rosette diameter, and flowering.

### Histochemical assay

Seedlings of promoter-reporter lines were incubated in GUS staining buffer, [100 mM sodium phosphate buffer (pH=7.2), 10 mM EDTA, 0.1% Triton X-100, 2 mM potassium ferricyanide, 2 mM potassium ferrocyanide and 1 mg ml–1 5-bromo-4-chloro-3-indolyl-β-D-glucuronide] at 37°C for 12 h. The chlorophyll was removed by incubation and multiple washes using 70% ethanol. The seedlings were observed under a Leica microscope (LAS version 4.12.0, Leica Microsystems) for the GUS staining.

### Confocal microscopy imaging

The fluorescence of GFP in Arabidopsis promoter-reporter lines seedlings (root) was observed using confocal microscopy (Zeiss LSM710, Zeiss LSM Image Examiner Version 4.2.0.121, CarlZeiss) at an excitation 495 nm/emission 545 nm.

### Expression analysis

Total RNA from seedlings, 30-day-old rosettes, of Arabidopsis was extracted using Spectrum Plant Total RNA kit (Sigma-Aldrich, USA). RNA’s quantity and quality were assessed using a spectrophotometer (NanoDrop, Thermo USA) and 1.2% agarose gel electrophoresis. DNA-free RNA (1 µg) was used to synthesize the first strand of cDNA using the Revert Aid First Strand cDNA Synthesis Kit (Fermentas, USA) as per manufacture instructions. Real-time PCR reactions were performed on an ABI 7500 Fast instrument (ABI Biosystems, USA). The expression was normalized using Tubulin and analysed through the comparative ΔΔ^CT^ method by Livak, and Schmittgen 2001. The primer sequences used for the expression analysis are listed in Supplemental Table S2.

### Western blot analysis

For protein extraction, whole seedlings (approximately 100 mg) were frozen in liquid nitrogen and ground in a protein extraction buffer containing 50 mM Tris (pH 7.5), 150 mM NaCl, 1 mM EDTA, 10% glycerol, 5 mM DTT, 1% protease inhibitor cocktail, and 0.1% Triton-X100. The resulting homogenate was centrifuged at 14,000 xg for 20 minutes at 4 °C in an eppendorf tube, and the supernatant was collected for further analysis. Protein concentrations were measured using the Bradford assay with a 5 μl aliquot from the supernatant (Bradford, 1976). Sodium dodecyl sulfate-polyacrylamide gel electrophoresis (SDS-PAGE) and western blot analysis were performed according to standardized methods (Sharma et al. 2020). For the western blot analysis, primary antibodies were diluted in TBS-T, specifically at a dilution of 1: 10000 (v/v) for anti-actin (Sigma-Aldrich) and 1:1000 (v/v) for anti-HY5. Western blot images were captured using Image Lab version 5.2.1 build 11 (Bio-Rad Laboratories).

### Quantification of total terpenoid

Total terpenoid content was estimated according to the previously described method by Ghorai et al. 2012. The plant material 500 mg 15-day-old seedlings were crushed in liquid N_2_ and homogenized in 3.5 ml of 95% methanol for 30 min in ice and then incubated at room temperature for 48 h in the dark with constant shaking. The mixture was centrifuged at 10,000xg for 15 min at 4°C, and 0.5 ml of supernatant was taken in a falcon tube. About 1.5 ml chloroform, 0.1 ml H_2_SO_4_, and 1.5 ml 95% methanol were added to it and incubated in ice for 1-2 h in the dark. The absorbance of the mixture was recorded at 538 nm. The total terpenoid concentration of the sample was calculated using the Linalool standard curve developed by using Linalool solution in methanol.

### GC-MS analysis

GC-MS analysis analysis, was carried out according to the previously described method by Michael et al. 2024. Fifteen-day-old seedlings were processed by crushing 300 mg of dried tissue into a fine powder using liquid N_2_, followed by homogenization with 12 ml of CHCl_3_/MeOH (2:1, v/v; Merck Millipore, USA). The extraction was carried out at 75°C for 60 minutes. Samples were dried using a Christ (Alpha 1-2 LD plus) lyophilizer and saponified at 90°C for 60 minutes in 2 ml of 10% (w/v) KOH in methanol with a rotator evaporator (Shimadzu QR 2005-S, Japan). After cooling, 1 ml of hexane and 1 ml of GC-grade H_2_O (Merck Millipore, USA) were added, and the mixture was shaken for 20 seconds on an orbital shaker (Geneilabs, India). Following centrifugation at 3,000 x g for 2 minutes, the hexane phase was transferred to a 2 ml glass vial (Agilent, USA) and re-extracted using another 1 ml of hexane. The combined organic phases were then evaporated to dryness. Next, 40 µl of N-methyl-N-(trimethylsilyl) trifluoroacetamide (MSTFA, Sigma Aldrich, USA) was added to the residue, shaken for 20 seconds, and transferred to a capped 2 ml auto-sampler glass vial (Agilent, USA) for incubation at room temperature for 5 minutes. Gas chromatography-mass spectrometry (GC-MS) analyses were performed using a Turbomass with Auto XL Perkin Elmer; Clarus 680GC instrument and a DB5MS column (30 m x 0.25 mm x 0.25 µm). The injector and mass selective detector (MSD) were set to 250°C. Analytes were separated at a flow rate of 1 ml/min with helium as the carrier gas and a 10:1 split ratio, following a thermal gradient from 100°C to 320°C. Analytes were fragmented in electron impact mode at 70 eV. Background signals were subtracted, and peaks were deconvoluted using software for identification based on mass fragmentation patterns compared against standards from the NIST and Wiley libraries. A blank injection was performed after each sample run. Five independent concentrations were used to create a standard curve for measuring phytosterol standards in the samples (Sigma-Aldrich, USA). The most abundant squalene ion has a mass-to-charge ratio (m/z) of 69.81, and squalene elutes from the column with a retention time of 11.88 minutes. The most abundant phytosterol ion has an m/z of 129, which is the parent ion for all phytosterols. The positive ion atmospheric pressure chemical ionization (APCI) mass spectrum for all phytosterols is presented in the right-hand panel. The retention times for the various phytosterols are as follows: Cholesterol at 13.72 minutes, Stigmasterol at 14.47 minutes, Beta-sitosterol at 13.15 minutes, Cycloartenol at 15.44 minutes, and Campesterol at 14.31 minutes.

### Exogenous Squalene assay

Synthetic squalene (Sigma-Aldrich, USA), which had a purity of 98%, was dissolved in hexane. The seedlings were grown on half-strength MS media having squalene concentrations ranging from 100-1000 ng/l. Hexane was included in the medium to serve as a reliable control, allowing for a clear comparison of the effects.

### Statistical analysis

The statistical tests and sample sizes, including biological replications, are detailed in the Fig. ligands. All statistical analyses were conducted using two-tailed Students’ t-tests with Graph Prism version 8.4.3 software. Each experiment was independently repeated at least three times, yielding consistent results.

### Accession numbers

Sequence data from this article can be found in the *Arabidopsis thaliana* Genome (Tair) under the following accession numbers: *AtHY5* (AT5G11260), *AtSQS1* (AT4G34640), *AtCHS* (AT5G13930), *AtSQE3* (AT4G37760) *AtCAS1* (AT2G07050) *AtDWF5* (AT1G50430).

## Acknowledgements

Author acknowledges Dr. Ranashekhar CH, Dr. Kapil Dev and Dr. Manju Singh from Central Instrumentation Facility, CSIR-CIMAP, for phytochemical analysis.

## Author contributions

P.K.T. conceived and designed the study. P.K.P, A.K., A.S. and N.S. participated in the execution of all the experiments. P.K.P and P.K.T. interpreted and discussed the data. P.K.P., N.S., A.S. and P.K.T. wrote the manuscript.

## Supplemental data

The following materials are available in the online version of this article.

**Supplemental Figure S1**. Schematic representation of major structural and biosynthetic genes of MVA and MEP pathways regulating terpenoid biosynthesis.

**Supplemental Figure S2**. Root phenotypic analysis of light-modulated plants in light and dark conditions.

**Supplemental Figure S3**. RT-qPCR expression analysis of AtCHS and AtHY5 is a positive control for light responsiveness.

**Supplemental Figure S4**. Representation of physical interaction of AtHY5 with AtSQS1 promoter in vitro through Y1H and EMSA.

**Supplemental Figure S5**. The ProAtSQS1 is negatively responsive towards the light in Arabidopsis WT, hy5-215 mutant, and HY5OX.

**Supplemental Figure S6**. Histochemical analysis of ProAtSQS1 in WT, hy5-215, and HY5OX in light and Dark.

**Supplemental Figure S7**. ProAtSQS1 exhibits negative responsiveness to light in *Arabidopsis thaliana*.

**Supplemental Figure S8**. Estimation of the total terpenoid in AtSQS1OX in Arabidopsis backgrounds.

**Supplemental Figure S9**. Phenotypic alterations on overexpression of AtSQS1 in WT and sqs1 backgrounds.

**Supplemental Figure S10**. Overexpression of AtSQS1 in hy5-215 and HY5OX backgrounds affects phenotypic and physiological parameters in *Arabidopsis thaliana*.

**Supplemental Figure S11**. Metabolic quantification of Phytosterols in Arabidopsis through GC-MS.

**Supplemental Figure S12**. Representation of CRISPR/Cas9-based editing in *SQS1^CR^/hy5-215* and SQS1OX/WT plants and modulation of squalene content due to resulting mutation.

**Supplemental Figure S13**. Expression analysis of terpenoid biosynthetic genes in *SQS1^CR^* /WT and *SQS1^CR^ /hy5-215* plants as compared to their respective controls.

**Supplemental Figure S14**. The representative image of the root length of light-modulated plants.

**Supplemental Figure S15**. Mutation in the AtSQS1 gene in *hy5-215* and WT backgrounds affects phenotypic and physiological parameters in *Arabidopsis thaliana*.

**Supplemental Figure S16**. Exogenous squalene is not able to restores seedling root growth in dark conditions in *SQS1^CR^*/WT.

**Supplemental Table S1**. Putative cis-acting light responsive elements (LRE) in *AtSQS1* Promoter.

**Supplemental Table S2**. List of primers and oligonucleotides used in this study.

## Data availability

All data generated or analyzed during this study are included in this published article (and its Supplemental information files).

